# Collection of Non Timber Forest Products (NTFPs) and their contribution to sustainable rural livelihoods in selected areas around Ayubia National Park, Pakistan

**DOI:** 10.1101/2023.11.22.568298

**Authors:** Mehwish Zaman, Asma Jabeen, Rabia Shabbir

**Affiliations:** Department of Environmental Sciences, Fatima Jinnah Women University, Rawalpindi, Pakistan

## Abstract

This study shows the Collection of Non Timber Forest Products (NTFPs) and their contribution to sustainable rural livelihoods in selected areas around Ayubia National Park, Pakistan. Forests are an important component for daily life of rural residents living near forested areas. A protected area is a geographically designated area for the purpose of biological conservation. Ayubia National Park (ANP) is a protected area in KPK, Pakistan that was designated with the purpose of management and preservation of nature. Non Timber Forest Products (NTFPs) have been defined as “All biological materials, excluding timber, which are extracted from forests for human use”. NTFPs play an important role in income generation for indigenous people. Malaach Village, Pasala Village and Khanuspur Town present around ANP were selected study areas. Purposive sampling was used for selection of study areas. Quantitative research was done. Field visits were done for Questionnaire survey. 50 questionnaires were filled from each selected village/town and results were analysed using SPSS. Cross tabulation was done and Chi Square test was applied. It was found that Wild Vegetables, Wild Fruits, Medicinal Plants and Mushrooms are collected from forests in and around ANP by indigenous people. Chi square results showed significant relationship between gender and collection of NTFPs. Poverty is found to be the main reason for collecting NTFPs. Contribution of NTFPs to annual income of people is mostly less than 20% but this contribution is also 21%-40% and even 41%-60% for few people. ANP should be managed properly to inhibit the collection of NTFPs within the park. Indigenous people should cooperate in sustainable collection and use of NTFPs. The role of NTFPs to livelihoods in and around ANP are clearly understood by study and will be useful in the awareness and implementation of the conservation practices in the Ayubia National Park.

## Introduction

Forests are an important component of the daily life of rural residents especially for those living near forested areas. Forests provide firewood, wood, forest soil, grasslands and raw materials to local residents [1–3]. Nearly 85% Pakistan’s forests are state owned, and the government is responsible for administration, conservation, and protection of those forests. Khyber Pakhtunkhwa Province contains over 40% of the entire countrywide forest area. The remaining forests are located in different provinces and areas. i.e., 14% are located in Baluchistan, 14.4% in Punjab, 9.4% in Sindh, 15.7% in Northern Areas and 6.5% is in Azad Jammu and Kashmir) [4]. Pakistan is one of the few areas on the planet with a unique biodiversity. It has a vast range of climate zones with a huge variety of plant species [5].

Pakistan’s has a great biodiversity resulting in distinctive floral compositions. Because of poverty, 80 percent of forest inhabitants rely on NTFPs in some form or another. The indigenous people depend on forests for the collection, handling, packing, drying, marketing, and use of various non-timber forest products. The forest dwellers rely on their indigenous knowledge about NTFPs. Several significant NTFPs present in Pakistan are morels, fruits as well as nuts, wild vegetable and herbs [6]. In Pakistan, forests have two distinct legal statuses: Forests owned by the state but not accessible to local communities. Those are known as state owned forests (generally including forests Reserved by state which are called Reserved Forests. These are Protected forests), and other forests called Guzara Forests or Communal forests which are given some level of access up to private ownership level . These kind of forest may be provided to either communities or individual persons. Reserved and Guzara both kind of forests are under Forest Department’s responsibility [7].

Parks and protected places are among the utmost effective tools for providing conservation to the biodiversity [8–11]. A protected area is a geographically defined area which is designated and managed to attain specific conservation objectives or an area (land or sea) which is specifically designated for the purpose of maintenance, protection and biological conservation [12].

Non Timber Forest Products (NTFPs) have been defined as “All biological materials, excluding timber, which are extracted from forests for human use” [13]. Non-timber forest products are biologically produced goods other than timber and fuelwood derived from forests and trees outside of forests. Non-Timber Forest Products play important part in income generation of local people. Non-timber forest products (NTFPs) are noteworthy for developing countries’ economies. NTFPs, include hundreds of traded and locally used biological forest products. These natural products derived from forest are progressively valued because of their contribution to rural livelihood, biodiversity conservation and export worth. NTFPs provide necessary food, nutrition and medicine for most of the local residents and families. These products are particularly important during the “hunger periods” in the agricultural cycle. These also prove to be helpful in seasonal variations. Poor indigenous people have more access on the forests than other important resources. Therefore, they are more dependent on these products for their livelihood [14]. The collections of wild non-timber forest products (NTFP) is a key source of income to millions of people throughout the World. Precise management methods are needed for a variety of NTFP species in order to gain the maximum collection benefits [15]. Local people often harvest NTFPs for trade as well as for household consumption. Some NTFPs, such as medicinal plants, can be symbolic and culturally significant, giving livelihood benefits through their social significance [16]. Asia is by far the largest producer and consumer of NTFPs in the World [17].

Beside an improved focus on the role of NTFPs in conservation, the past some decades have also seen a growing attention and effort in establishing protected areas (PAs). Protected areas worldwide are a key approach to address the problem of enormous forest and biodiversity damage [18; 19]. Ayubia National Park in Abbotabad was designated with the purpose of preservation of nature. The goal of Ayubia National Park is to preserve nature and natural processes in a viable representative area of the forests in Gallies. Summer season is moderate in this area and winter season is really harsh with considerable snowfall at high elevations. The amount of rainfall varies each year in this area. There is a high amount of precipitation in the area. The month of July–August receives 60% of the average precipitation, with the remaining 40% unequally divided between September and June. Poverty is dominant in the area and is causing threatening proportions with passing time. The economy of Abbotabad is depending comprehensively on natural resources. The nourishment and sustenance by agriculture is a major part of this dependence [20].

NTFPs collection is done from the Reserved as well as from Guzara forests. A diversity of wild vegetables are present in the Ayubia National Park. The surrounding populations of the Park collect numerous types of wild vegetables. These NTFPs are frequently gathered among the month of April and the end of June [21]. The main vegetables collected are Kunji (*Dryopteris stewartii*), Mushkana (*Nepeta laevigata*), Kandhor (*Dryopteris blanfordii*), Mirchi (*Solanum nigra*), and Tandi (*Dipsacus inermis*). The parts collected are young leaves for all species [22].

## Materials and Methods

Questionnaire survey was done and data was collected for analysis purpose. This was a survey based research in which questionnaire survey was done by collecting information from indigenous people living around selected areas of Ayubia National Park, Pakistan. This research work was approved by Advanced Studies and Research Board (AS & RB) of Fatima Jinnah Women University, Rawalpindi, Pakistan. This study does not involved samples from human participants so there was not any need of approval of this study from the ethics committee.

Before starting the survey work, villages were selected for data collection. There are total eight villages around Ayubia National Park [23] and three small town Thandiani, Nathiagali and Khanuspur [24]. Malaach, Pasala, Moorti, Kuzagali, Darwaza, Riala, Lahur Kas, and Khunkhurd are the main villages around Ayubia National Park area [25]. Two villages (Malaach and Pasala) and a town (Khanuspur) surrounding Ayubia National Park were the selected study areas of the following research. Questionnaire survey was done from the selected study areas for studying the contribution of Non Timber Forest Products to livelihoods of indigenous people in the selected villages /towns.

Before taking information from the indigenous people informed consent was taken from them. The research participants were informed about the details of the research work and verbal consent was taken from them. The consent was documented by audio recording. Most of the people living around the villages of Ayubia National Park had less or no education so they did not provided any written consent and they only verbally explained that they are ready to provide the required information. There were not any children involved in the study so there was not any need of approval from their parents about information collection. There was not any need of approval of consent from ethics committee because the following study does not involve any medical records or samples.

The survey and field work was done during May till August in 2019. Questionnaires were filled by doing extensive field visits. Purposive Sampling was used for the selection of villages and houses from those villages/town were selected randomly. Permission was taken from the Forest and the Wildlife Department of KPK about the field visits and for gathering information from the indigenous people living in selected villages around Ayubia National Park. Ayubia National Park is managed by both Wildlife Department and forest of the Khyber Pakhtunkhwa Province, Pakistan. The Forest and Wildlife officers verbally provided permission for taking information related to research from selected areas around Ayubia National Park, Pakistan.

### Study Area

Ayubia National Park is a 3312-hectare protected area in Abbottabad, KPK, and Northern Pakistan. In 1984, it was designated as a national park. Ayubia was named upon Pakistan’s 2nd President, Muhammad Ayub Khan (1958-1969). The elevation of this area is 8000 feet above sea level [26]. Ayubia National Park is located in Gallies Forest division of Abbotabad. The park is situated on array of hills in a row north to south near Abbottabad and the northwestern side of Murree. Headquarter of the park is situated at Dungagali. Dungagali is located at 43 kilometers from Abbottabad as well as 30 kilometers from Murree [27; 23]. Around 50,000 residents are living nearby the Park in different villages [28]. The park has been designated as part of the Galliat Reserved Forests [29]. It lies among 34°-1’ and 34°-3.8’ north latitude and 73°-22.8’ to 73°-27.1’ east longitude. It was originally designated in 1984 and covered an area of 1684 hectares. The area of Ayubia National Park was expanded under the KPK Wildlife Act of 1975. The park’s area was increased and it was more than doubled to 3,312 hectares in March 1998 [2].

In order to analyze the contribution of NTFPs to livelihoods, two villages and a town were selected around Ayubia National Park. These are **Malaach**, **Pasala** and **Khanuspur**. Selected villages/ town around Ayubia National Park were the study areas of this research (Fig 1). These villages and town were selected purposively for this research. Malaach and Pasala villages are located in the reserved forests and these were purposively selected in order to know the contribution of Non Timber Forest Products to livelihoods of the local people living in these villages.

**Fig 1.**
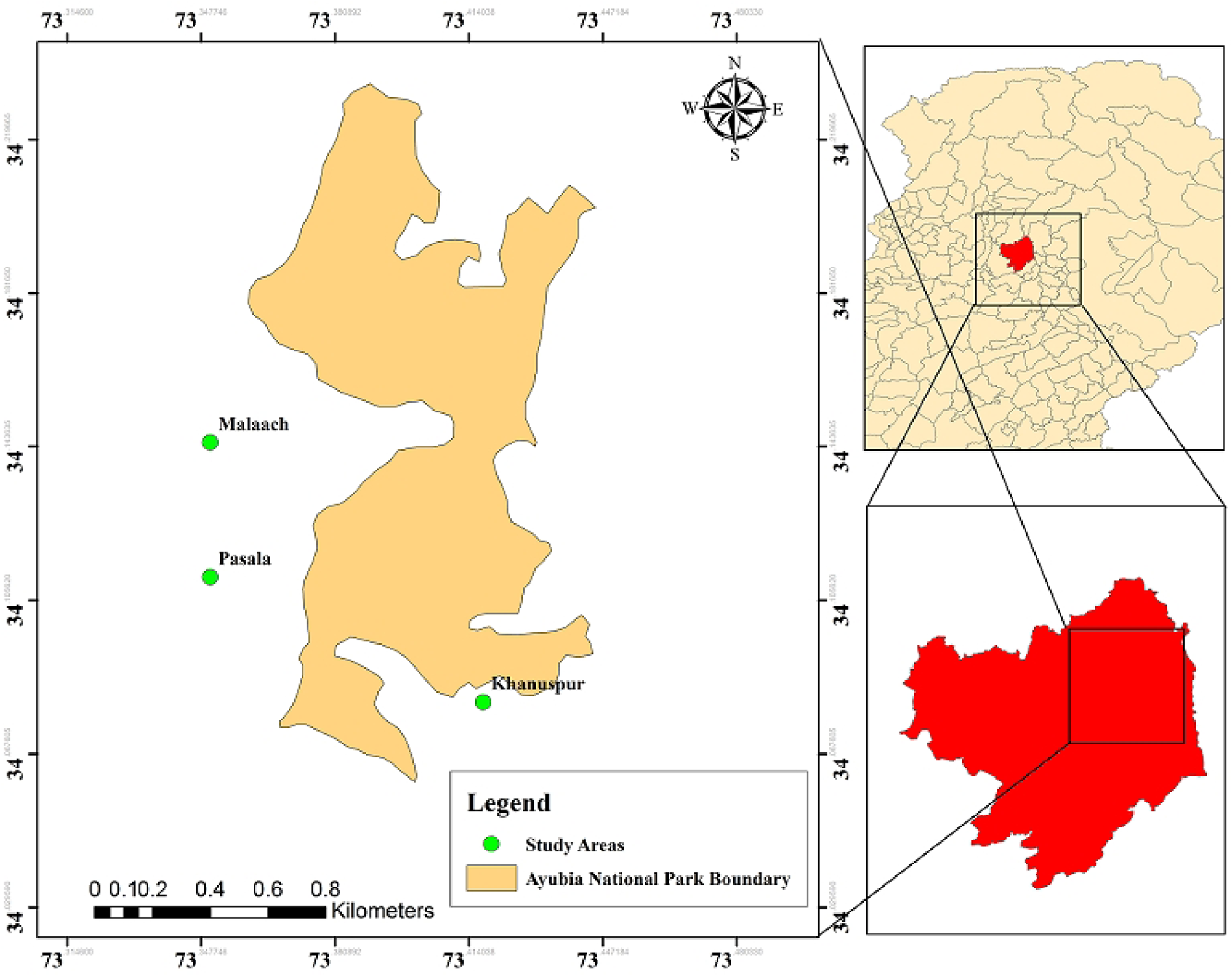
Map showing selected study areas around Ayubia National Park, Pakistan.

### Sampling Method

Purposive sampling was used for the selection of villages around ANP for research. Purposive sampling is also known as judgement sampling. It is the purposeful or intentional selection of a participant based on the participant’s characteristics or potentials. In purposive sampling, it is absolutely the choice of the researcher to decide that is necessary to be known. Purposive sampling involves identifying and selecting individuals, areas, or groups of individuals who are knowledgeable and capable in a topic of interest [30]. In this research the villages and town were selected purposively on the basis of the judgment made by contacting people about the collection and usage of NTFPs. During the selection of villages and town, the villages with the most local people households dependent on NTFPs were given priority [13].

### Household Level Analysis (Questionnaire survey)

Questionnaires were filled by visiting mainly the houses in the selected villages and town. A combination of random and purposive sampling technique were employed in selecting the respondents [31]. The houses from those villages were randomly selected. This survey included the household level analysis. For the household analysis the structured quantitative survey was used [32]. In each village/town selected, 50 respondents/houses were randomly picked for the data collection as the exact population of each village was not known. Hence, 50 questionnaires were filled from each village/town. The questionnaire was designed in such a manner that it contained all the necessary questions related to importance and contribution of the non timber forest products in the livelihoods of Ayubia National Park. For the analysis of the questionnaire survey, Statistical Package for Social Sciences (SPSS) and EXCEL software packages were used. Data was entered on Excel and then SPSS software version 23 was used for analysis. Descriptive statistics was used to analyze the data [31]. Clustered bar charts were made from the collected data of questionnaire. Cross tabulation was done and chi square was applied.

## Results

Clustered bar charts were made related to some of the important results found in the following study. Clustered bar charts were made about gender and reason for collection of NTFPs, and between gender and contribution of NTFPs to annual income in the three selected areas (villages/towns). Cross tabulation tables were made showing the observed and expected counts and chi square test was done to find the relationship between variables.

### Poverty as reason for collection of NTFPs

Reason for collection of NTFPs for maximum respondents in Malaach is poverty. According to 23 respondents (20 females and 3 males) the main reason for collection of NTFPs in Malaach is poverty. According to 9 respondents (6 Females and 3 males) the reason for collection of NTFPs is Business i.e, they sell the collected NTFPs for income generation. Personal interest is the main reason for collection of NTFPs according to 7 respondents (3 females and 4 males). According to 11 respondent (1 female and 10 males) from Malaach there is not any reason for collection of NTFPs because they do not collect NTFPs (Fig 2 a).

**Fig 2.**
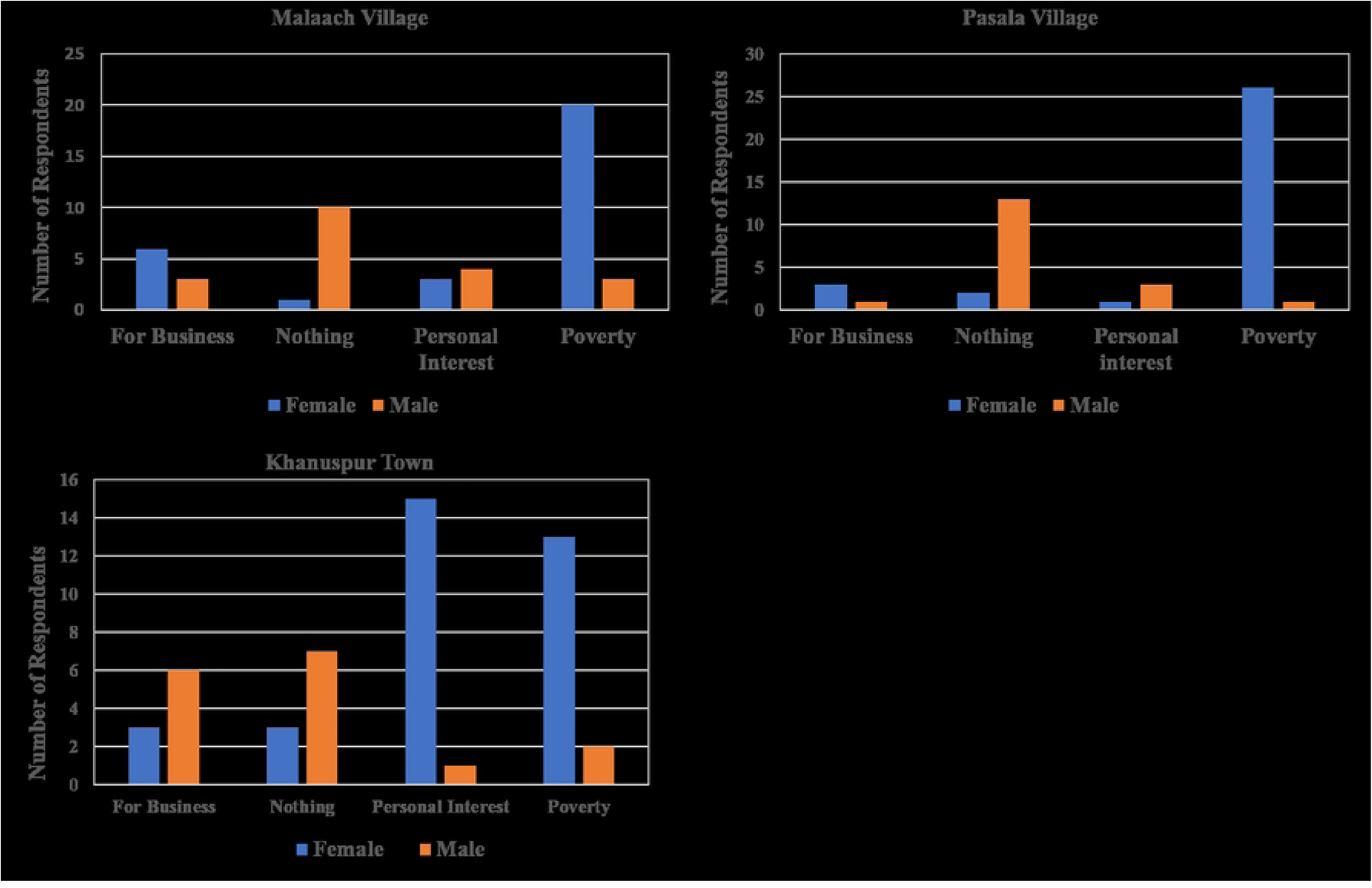
Reason for collection of NTFPs by gender in study areas.

Reason for collection of NTFPs in Pasala also is poverty. According to 27 respondents (26 Females and 1 male) main reason for collection of NTFPs in Pasala is poverty. According to 4 respondents (3 females and 1 male) reason for collection is business. According to 4 respondents (1 female and 3 males) the reason for collection of NTFPs in Pasala is personal interest. 15 respondents (2 females and 13 males do not collect NTFPs in Pasala so there is not any reason for collection of NTFPs according to them (Fig 2 b). Reason for collection of NTFPs in Khanuspur town is mainly personal interest mostly and also poverty. According to 16 respondents (15 females and 1 male), reason for collection of NTFPs in Khanuspur is personal interest. According to 15 respondents (13 females and 2 males) reason for collection of NTFPs is poverty. According to 9 (3 females and 6 males) respondents the reason for collection is business. 10 respondents (3 females and 7 males) do not collect NTFPS in Khanuspur town, so there is not any reason for collection (Fig 2 c). A variety of Non Timber Forest Products (Wild Vegetables, Wild Fruits, Mushroom and Medicinal Plants) are collected by indigenous people living in selected study areas around Ayubia National Park (Lists in SI File). NTFPs are contributing to livelihoods of indigenous people.

Table 1, Table 2 and Table 3 show the Reason for collection of NTFPs by gender in Malaach village, Pasala village and Khanuspur town.

**Table 1.**
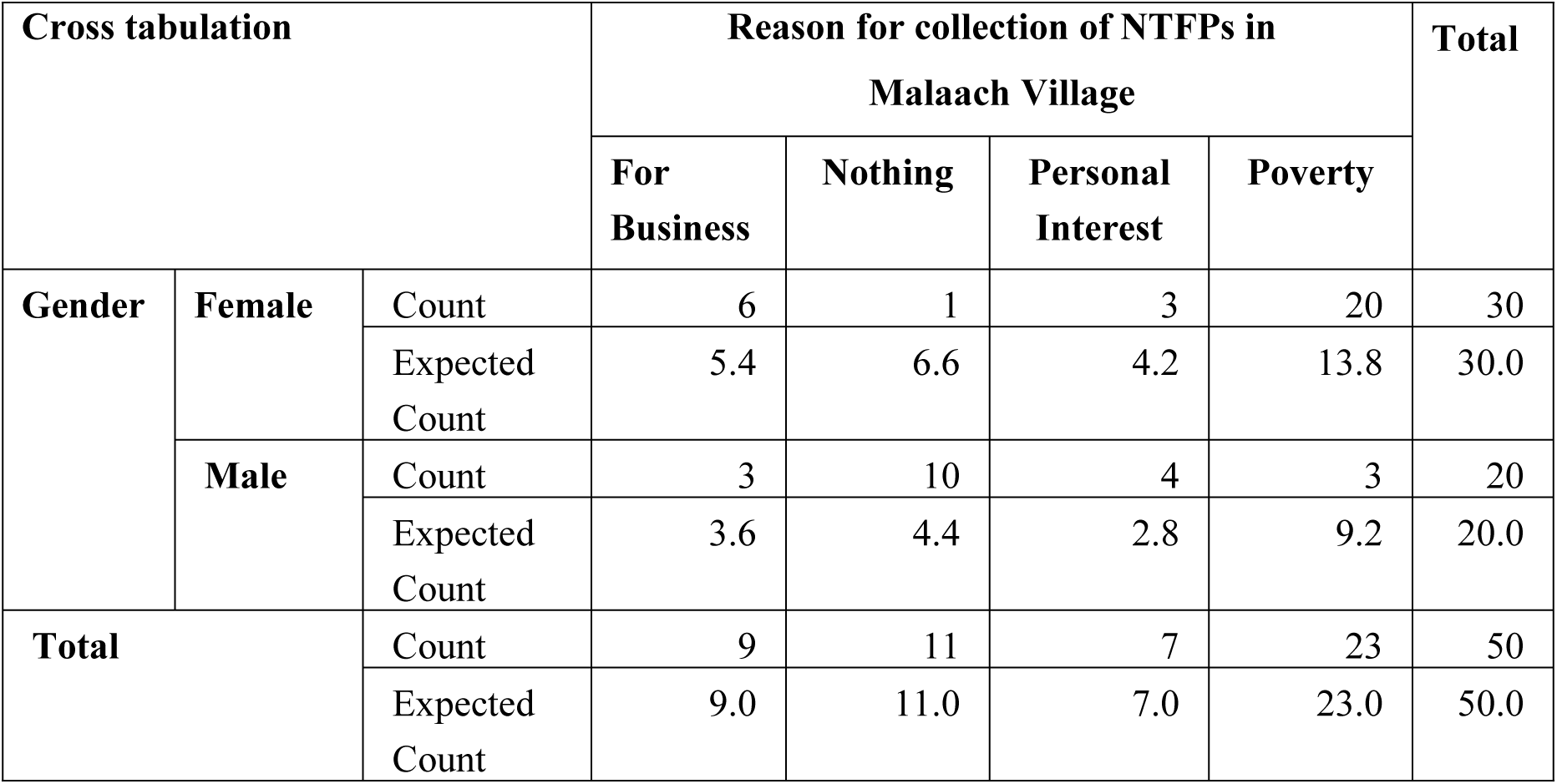
Gender and Reason for collection of NTFPs in Malaach Village.

**Table 2.** Gender and Reason for collection of NTFPs in Pasala Village.

| Cross Tabulation |  |  | Reason for collection of NTFPs in Pasala Village |  |  |  | Total |
| --- | --- | --- | --- | --- | --- | --- | --- |
|  |  |  | For Business | Nothing | Personal interest | Poverty |  |
| Gender | Female | Count | 3 | 2 | 1 | 26 | 32 |
|  |  | Expected Count | 2.6 | 9.6 | 2.6 | 17.3 | 32.0 |
|  | Male | Count | 1 | 13 | 3 | 1 | 18 |
|  |  | Expected Count | 1.4 | 5.4 | 1.4 | 9.7 | 18.0 |
| Total |  | Count | 4 | 15 | 4 | 27 | 50 |
|  |  | Expected Count | 4.0 | 15.0 | 4.0 | 27.0 | 50.0 |

**Table 3.**
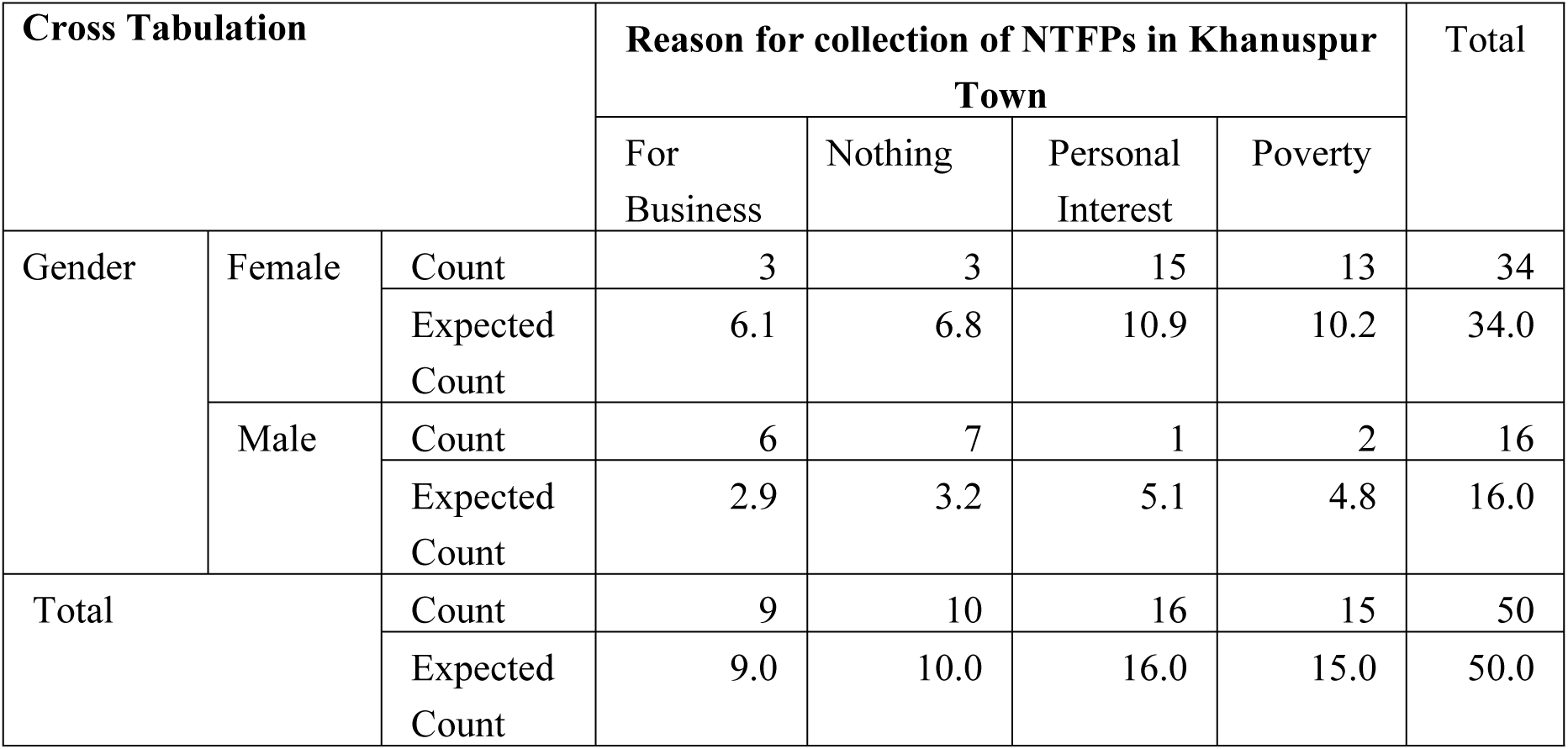
Gender and Reason for collection of NTFPs in Khanuspur Town.

| Cross Tabulation |  |  | Reason for collection of NTFPs in Khanuspur Town |  |  |  | Total |
| --- | --- | --- | --- | --- | --- | --- | --- |
|  |  |  | For Business | Nothing | Personal Interest | Poverty |  |
| Gender | Female | Count | 3 | 3 | 15 | 13 | 34 |
|  |  | Expected Count | 6.1 | 6.8 | 10.9 | 10.2 | 34.0 |
|  | Male | Count | 6 | 7 | 1 | 2 | 16 |
|  |  | Expected Count | 2.9 | 3.2 | 5.1 | 4.8 | 16.0 |
| Total |  | Count | 9 | 10 | 16 | 15 | 50 |
|  |  | Expected Count | 9.0 | 10.0 | 16.0 | 15.0 | 50.0 |

The observed count for Gender and reason for collection of NTFPs in Malaach village shows that 6 females and 3 males collect NTFPs for business purpose, 3 females and 3 males collect NTFPs because of personal interest, and 20 females and 3 males collect NTFPs because of poverty in Malaach Village. There is not any reason of collection for 1 women as she does not collect NTFPs. The observed count for Gender and reason for collection of NTFPs in Pasala Village shows that 3 females and 1 male collect NTFPs for business purpose, 1 female and 3 males collect NTFPs because of personal interest and 26 females and 1 male collect NTFPs because of poverty in Pasala Village. There is not any reason of collection for 2 females and 13 males as they do not collect NTFPs. The observed count for Gender and reason for collection of NTFPs in Khanuspur town shows that 3 females and 6 males collect NTFPs for business purpose, 15 females and 1 male collect NTFPs because of personal interest and 13 females and 2 males collect NTFPs because of poverty in Khanuspur Village. There is not any reason of collection for 3 females and 7 males as they do not collect NTFPs.

**Table 4.** Chi Square results for Gender and Reason for collection of NTFPs in selected study areas.

| Chi Square Tests | Malaach Village |  |  | Pasala Village |  |  | Khanuspur Town |  |  |
| --- | --- | --- | --- | --- | --- | --- | --- | --- | --- |
|  | Value | df | Asymptotic Significance (2-sided) | Value | df | Asymptotic Significance (2-sided) | Value | df | Asymptotic Significance (2-sided) |
| Pearson Chi-Square | 19.866<sup>a</sup> | 3 | .000 | 31.787<sup>a</sup> | 3 | .000 | 18.884<sup>a</sup> | 3 | .000 |
| Likelihood Ratio | 21.769 | 3 | .000 | 36.010 | 3 | .000 | 19.751 | 3 | .000 |
| N of Valid Cases | 50 |  |  | 50 |  |  | 50 |  |  |
|  | 4 cells (50.0%) have expected count less than 5. The minimum expected count is 2.80. |  |  | 4 cells (50.0%) have expected count less than 5. The minimum expected count is 1.44. |  |  | 3 cells (37.5%) have expected count less than 5. The minimum expected count is 2.88. |  |  |

The Chi Square result in Table 4 show the significant relationship between the variables gender and reason for collection in all three selected study areas. The value of Chi square is .000 for all three study areas. It is concluded that the variables are associated with eachother.

### Contribution of NTFPs to Annual Income of Indigenous People

NTFPs contribute to the annual income of indigenous people. According to 18 respondents (12 females and 6 males) the contribution of NTFPs to annual income is less than 20% in Malaach. According to 7 respondents (4 females and 3 males), the contribution of NTFPs to annual income in Malaach Village is 21%-40%. According to 2 female respondents, the contribution of NTFPs to annual income in Malaach village is 41%-60%. According to 23 respondents (12 females and 11 males) from Malaach village, there is not any contribution of NTFPs to income (Fig 3 a).

**Fig 3.**
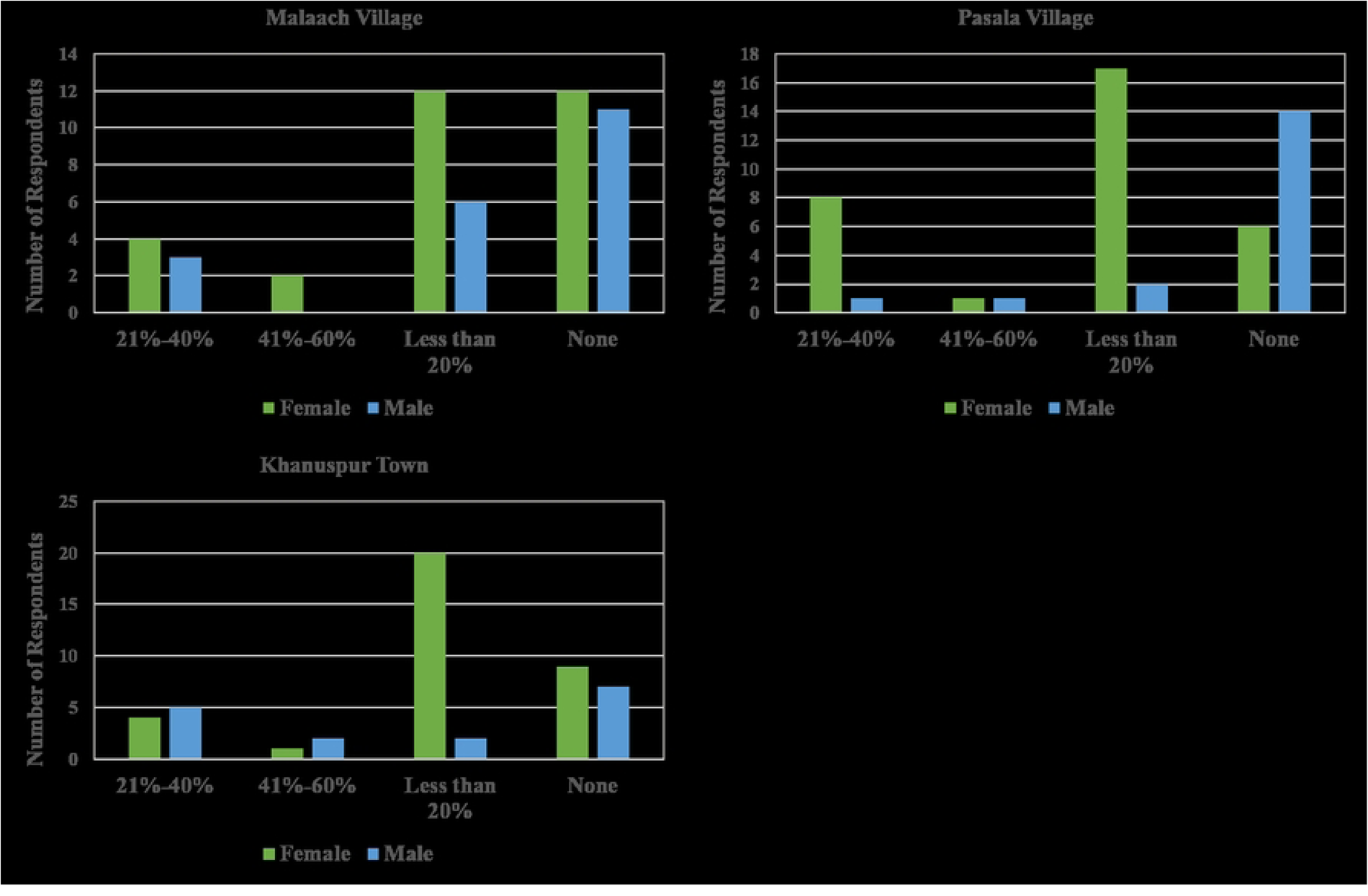
Contribution of NTFPs to annual income of indigenous people in study areas.

According to 19 respondents (17 females and 2 males), the contribution of NTFPs to annual income in Pasala village is less than 20%. According to 9 respondents (8 females and 1 male) the contribution of NTFP to annual income is 21%-40%. According to 2 respondents (1 male and 1 female) the contribution of NTFPs to annual income is 41%-60%. According to 20 respondents (6 females and 14 males) in Pasala there is not any contribution of NTFPs to annual income (Fig 3 b).

For maximum respondents of Khanuspur, contribution of NTFPs to annual income is less than 20%. According to 22 respondents (20 females and 2 males) the contribution of NTFPs to annual income in Khanuspur Town is less than 20%. According to 9 respondents (4 females and 5 males) of Khanuspur, the contribution of NTFPs to annual income is 21%-40%. According to 3 respondents (1 female and 2 males), the contribution of NTFPs to annual income is 41%-60%. There is not any contribution to annual income according to 16 respondents (9 females and 7 males) from Khanuspur town (Fig 3c).

Table 5, Table 6 and Table 7 show Contribution of NTFPs to annual income of males and females in Malaach, Pasala and Khanuspur.

**Table 5.** Gender and Contribution of NTFPs to Annual Income in Malaach Village.

| Cross tabulation |  |  | Contribution of NTFPs to Annual Income in Malaach Village |  |  |  | Total |
| --- | --- | --- | --- | --- | --- | --- | --- |
|  |  |  | 21%-40% | 41%-60% | Less than 20% | None |  |
| Gender | Female | Count | 4 | 2 | 12 | 12 | 30 |
|  |  | Expected Count | 4.2 | 1.2 | 10.8 | 13.8 | 30.0 |
|  | Male | Count | 3 | 0 | 6 | 11 | 20 |
|  |  | Expected Count | 2.8 | .8 | 7.2 | 9.2 | 20.0 |
| Total |  | Count | 7 | 2 | 18 | 23 | 50 |
|  |  | Expected Count | 7.0 | 2.0 | 18.0 | 23.0 | 50.0 |

**Table 6.** Gender and Contribution of NTFPs to Annual Income in Pasala Village.

| Cross Tabulation |  |  | Contribution of NTFPs to Annual Income in<br>Pasala Village |  |  |  | Total |
| --- | --- | --- | --- | --- | --- | --- | --- |
|  |  |  | 21%-40% | 41%-60% | Less than<br>20% | None |  |
| Gender | Female | Count | 8 | 1 | 17 | 6 | 32 |
|  |  | Expected<br>Count | 5.8 | 1.3 | 12.2 | 12.8 | 32.0 |
|  | Male | Count | 1 | 1 | 2 | 14 | 18 |
|  |  | Expected<br>Count | 3.2 | .7 | 6.8 | 7.2 | 18.0 |
| Total |  | Count | 9 | 2 | 19 | 20 | 50 |
|  |  | Expected<br>Count | 9.0 | 2.0 | 19.0 | 20.0 | 50.0 |

**Table 7.** Gender and Contribution of NTFPs to Annual Income in Khanuspur Town.

| Cross Tabulation |  |  | Contribution of NTFPs to Annual Income in<br>Khanuspur Town |  |  |  | Total |
| --- | --- | --- | --- | --- | --- | --- | --- |
|  |  |  | 21%-40% | 41%-60% | Less than<br>20% | None |  |
| Gender | Female | Count | 4 | 1 | 20 | 9 | 34 |
|  |  | Expected<br>Count | 6.1 | 2.0 | 15.0 | 10.9 | 34.0 |
|  | Male | Count | 5 | 2 | 2 | 7 | 16 |
|  |  | Expected<br>Count | 2.9 | 1.0 | 7.0 | 5.1 | 16.0 |
| Total |  | Count | 9 | 3 | 22 | 16 | 50 |
|  |  | Expected<br>Count | 9.0 | 3.0 | 22.0 | 16.0 | 50.0 |

The observed count for Gender and contribution of NTFPs to Annual Income in Malaach Village shows that 4 women have annual income 21%-40%, 2 women have annual income 41%-60%, 12 women have less than 20% income and 12 women have none income. 3 out of 20 males have annual income 21%-40%, none of the men have annual income 41%-60%, 6 men have annual income less than 20% and 11 men have none income in Malaach village (Table 5). The observed count for Gender and Contribution of NTFPs to Annual Income in Pasala Village shows that 8 females and 1 male have 21%-40% annual income, 1 female and 1 male have 41-60%, 17 females and 2 males have less than 20%, and 9 females and 14 males have no income because of NTFPs (Table 6). The observed count for Gender and contribution to annual income in Khanuspur Town shows 4 women and 5 males have annual income of 21%-40%, 1 female and 2 males have 41%-60% annual income because of NTFPs, 20 females and 2 males have annual income less than 20% and 9 females and 7 males have no income because of NTFPs (Table 7).

The Chi Square results in Table 8 show the value of Chi square is .000 for Pasala Village which show significant relationship between the variables Gender and Contribution of NTFPs to annual income. The value of chi square is 0.517 for Malaach village which means there is not significant relationship between variables. The variables are not associated with each other. Contribution of NTFPs to annual income is less in Malaach village and that contribution is more in Pasala and Khanuspur village. The Chi square result of Khanuspur town shows that there is significant relationship between the variables gender and Contribution of NTFPs to annual income in Khanuspur Town. The Chi square result of Khanuspur town is .016 which is less than 0.05 so there is a significant relationship between the variables.

**Table 8.**
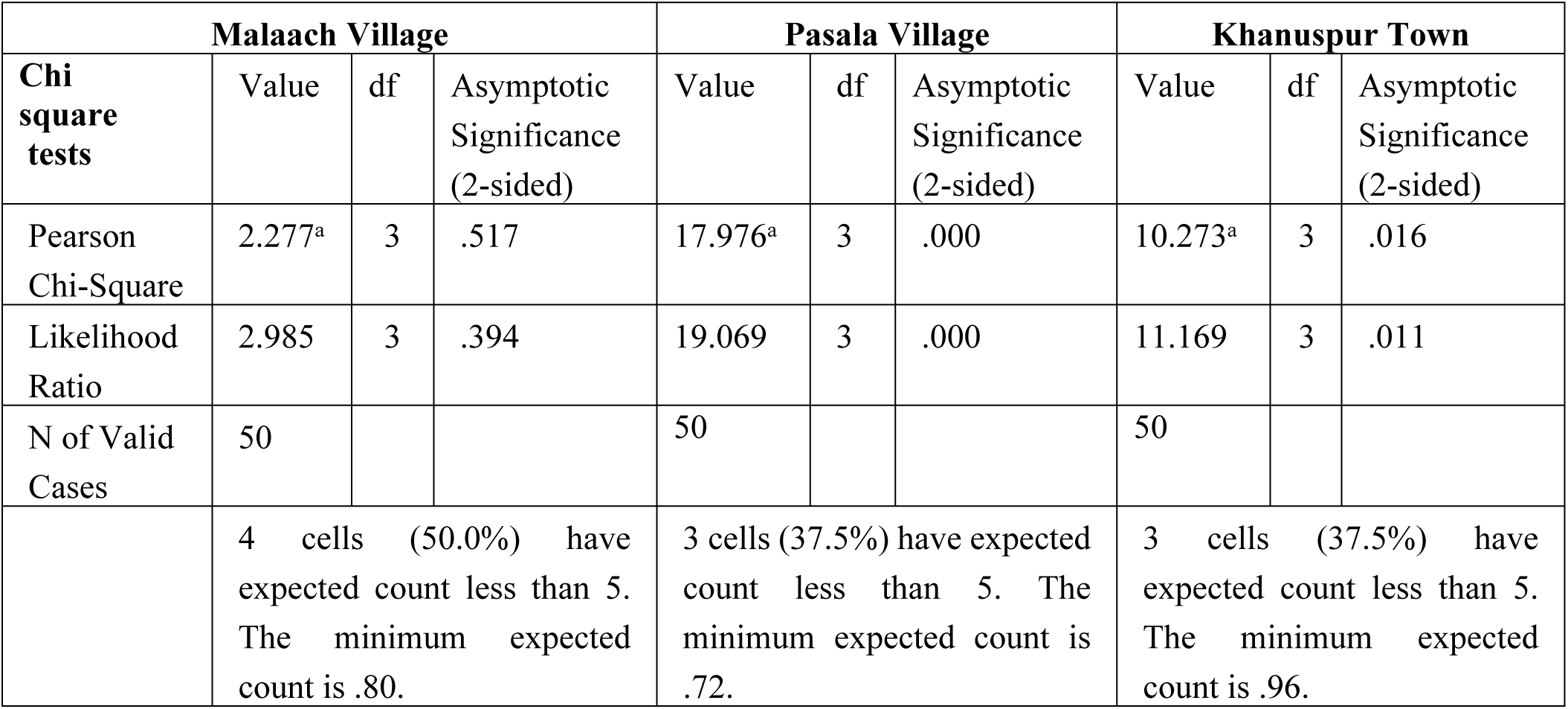
Chi Square results for Gender and Contribution of NTFPs to Annual Income in selected study areas.

## Discussion

In the following research it was found that indigenous people are reliant on the Non Timber forest products for sustaining their livelihoods from Malaach, Pasala and Khanuspur. The quantitative results showed the dependency of people on Non timber forest products. NTFPs play a variety of roles depending on the economic and cultural situations in which they are used. The value of NTFPs differs according to the economic perspectives and cultural backgrounds. In developed countries, NTFPs are often used for recreational and cultural purposes, conservation of biodiversity and rural economic growth. They’re primarily used for subsistence and income generating in poor countries, particularly in Africa and Asia. Challenges to more than 99% of the world’s protected areas include poaching, encroachment from farming, and unsustainable collection of non-timber forest products (NTFPs) [33; 34]. NTFPs are thus seen as a security net in underdeveloped nations, filling in the gap caused by a lack of agricultural output or other emergency situations [35–37]. As Pakistan is also a developing country and the study area of this research are villages so indigenous people of those areas are dependent on NTFPs.

*Taxus wallichiana* (Yew), *Valeriana jatamansi* (Mushk bala), *Bergenia ligulata* (Zakham-e-Hayat), *Podophyllum emodi* (Bankakri) and *Skimmia laureola* (Nair) are several main collected medicinal plants from ANP [27]. A number of wild vegetables grow in and around Ayubia National Park. The surrounding and nearby populations of the park collect and eat several kinds of wild vegetables. Wild vegetable varieties Kunji saag (*Dryopteris stewartii),* Mushkana saag *(Nepeta laevigata),* Kandhor saag *(Dryopteris blanfordii),* Mirchi *(Solanum nigra), Tandi (Dispacus inermis* are gathered from ANP [24; 38]. The ANP yields a large number of therapeutic plants and a lot of these medicinal herbs are gathered from the ANP [22]. Women and children are mostly gatherers of these plants [38].

Forests are an important component of an economy’s natural capital. Most importantly, NTFPs make a considerable contribution to rural household income in many nations around the world. In Central and West Africa, the portion of household income derived from NTFPs profits is often equal to or more than the minimum wage for school teachers. The lowest households’ dependence on forest revenue is more significant (as a percentage of their total annual household income [39]. According to a study of Indonesia Natural forest revenue was quite high for indigenous households. The income of indigenous people from forests was 36%. This is probable because only NTFPs that don’t require expensive equipment are extracted [40]. According to following research most of the people of Malaach, Pasala and Khanuspur collect the NTFPs due to poverty as shown in Tables 1, Table 2 and Table 3. From Malaach village, 23 respondents (46%) have been collecting NTFPs because of poverty. From Pasala, 27 respondent (54%) have been collecting NTFPs because of poverty and from khanuspur 15 respondents (30%) have been collecting NTFPs because of poverty. Khanuspur respondents collect mostly because of personal interest also.

NTFPs have the ability to reduce seasonal and long-term hunger as well as food shortages. Indigenous people have knowledge about the availability of forest resources. Indigenous people occasionally generate income from these sources. This as a result which gives an alternate livelihood source for them. Non-timber forest products (NTFPs) are primarily collected, handled, and sold by women. Children and women both are engaged in the collecting and sale of various NTFPs, but women’s participation is higher as compared to men [41].

Indigenous people in Pakistan’s mountainous areas collect around 600 medicinal plant species (around 10% of the total documented plant species) to support their livelihoods (Shinwari 2010). Morel gathering and processing are significant economic activity throughout Pakistan’s Hindu Kush Himalayas (Khyber Pakhtunkhwa province, KPK). Over 5,000 local households are involved in collection of morels in KPK [42; 43].

The forests around Malaach and Pasala village are mostly reserved forests and local people of Malaach and Pasala collect NTFPs from those reserved forest of ANP. Local populations harvest a variety of medicinal plants from the Ayubia National Park [44]. Mushrooms are among the most expensive Non-Timber Forest Products (NTFPs) in the region, and as a result, they attract a large number of collectors. NTFPs are increasingly being acknowledged for their significance to the livelihoods of a huge number of people in developing nations. For millions of people, there is a tremendous job opportunity by NTFPs [14]. The present research showed that NTFPS are an important source of annual income for indigenous people of Malaach, Pasala and Khanuspur. Contribution of NTFPs to annual income of people is mostly less than 20% but this contribution is also 21%-40% and even 41%-60% for few people. It is stated that 15-30% of the overall income of poor households in developing countries is by medicinal plants [45].

Non-timber forest products play an important role in rural income generation through protected forests. These NTFPs have not been well recognized and promoted for income generation for a sustainable or long term basis. In self-financed and self-managed development in rural areas, income creation through NTFPs and other resources can play a key role. Many NTFP collection procedures include the less-harmful harvesting of annual renewable plant parts, which provides a profitable revenue stream for improving community income [14].

In present research the contribution of NTFPs to annual income in Malaach village for 18 respondents (36%) is less than 20%. For 7 respondents (14%) the contribution of NTFPs to annual income is 21%-40% and there is 41%-60% contribution of NTFPs to annual income for 2 respondents (4%) of Malaach. For maximum respondents the contribution to annual income in Malaach is less than 20%. For Pasala village respondents also NTFPs are contributing to annual income. There is less than 20% contribution of NTFPs to annual income for 19 Pasala respondents (38%) and 21%-40% contribution to annual income for 9 respondents (18%) of Pasala. There is 41%-60% contribution to annual income by NTFPs for 2 Pasala respondents (4%). For Khanuspur town also there is contribution of NTFPs to annual income. For 19 respondents (38%) of Khanuspur the contribution to annual income is less than 20% and for 9 respondents (18%) this contribution is 21%-40%. For 6 respondents (12%) of Khanuspur the contribution of NTFPs to annual income is 41%-60%. It is stated that 15-30% of the overall income of poor households in developing countries is by medicinal plants [45].

Rural areas are home to about 75% of the poor in developing countries [46]. According to the Global multidimensional poverty index MPI 2014, 85% of poor people are living in rural areas according to the Global multidimensional poverty index MPI 2014, [47]. Their survival is typically reliant on agriculture, cattle and gathering of forest products [48]. The world is struggling with uncountable difficulties, including rising poverty conditions in numerous countries; particularly among the forest-dependent people. The majority of these settlements are located in distant places with few services and necessities. As a result, small communities frequently become overly reliant on the natural resources in their immediate vicinity. As a result, forest products, especially non timber forest products (NTFPs), have become significant and vital basis of income for the majority of forest-dependent people [49]. NTFPs significantly contribute to local livelihoods and even offer a pathway out of poverty [50]. Unsustainable collection of NTFPs such as medicinal plants for the purpose of sale, and unsustainable collection of fuelwood are all threatening biodiversity. The social, economic, and political circumstances are required for sustainable extraction and use of NTFPs [51; 52].

## Conclusion

It was concluded that NTFPs are playing a significant role in the livelihood of people living in selected villages/town around Ayubia National Park. Ayubia National Park is a protected area in KPK. There are eight villages around Ayubia National Park. There are three towns (Nathiagali, Khanuspur and Thandiani) also around ANP. It was concluded that in all the selected villages and town (Malaach, Pasala and Khanuspur), Non Timber Forest Products are collected and these are contributing to the livelihoods of people in many ways and people also generate income from those NTFPs. Wild Vegetables, Wild Fruits, Medicinal plants and Mushrooms are all considered important for the rural people. These are a source of food during poverty. There is a significant relationship between gender and collection of NTFPs in all the selected study areas. NTFPs are used in everyday life of the indigenous people. Contribution of NTFPs to annual income for maximum respondents in the selected villages and town is less than 20% and 21%-40% also. Most important NTFPs for livelihoods of people as according to these villages is Medicinal plants and wild vegetables mostly in Malaach village. Wild vegetables and mushrooms mostly in Pasala village while Medicinal Plants and mushrooms in Khanuspur town. It is concluded that there is also need of conservation for the NTFPs in and around the areas of Ayubia National Park.

## Acknowledgements

The authors are highly grateful to the indigenous people living in the selected areas around Ayubia National Park for providing useful information for questionnaire survey about Contribution of Non Timber Forest to their livelihoods.

